# The HIV-1 restriction factor RPRD2 does not inhibit transcription of HIV-1 or endogenous retroelements

**DOI:** 10.64898/2026.07.21.739879

**Authors:** Kathryn A. Jackson-Jones, Hui Xin Loh, Arda Tarcan, Rozeena Arif, Katie Duckett, Rebecca M. Fu, Connor Hird, Cameron Ferguson, Alfredo Castillo, Richard D. Sloan

## Abstract

A vital question in transcription regulation is how and to what extent transcription of different DNA species, including viral DNA, episomal DNA and endogenous retroelements, is differentially regulated to transcription of host genes. RPRD2 has previously been characterized as an HIV restriction factor that blocks reverse transcription. However, RPRD2 has also been characterized as a regulator of global transcription of host genes. Therefore, we hypothesized that RPRD2 may also regulate nascent transcription from HIV provirus and endogenous retroelements. First, we used a combination of experimental and computational methods to characterize the binding of RPRD2 to RNA and DNA:RNA hybrids. Using immunoprecipitation, we identified that RPRD2 interacts with both the transcription regulator PAF1 and the HUSH complex member TASOR, independently. Immunofluorescence revealed that GFP-RPRD2 localizes to foci in the nucleus, and these foci overlap with nuclear speckles. To measure the effect of RPRD2 on transcription, we used plasmid-borne HIV LTR-driven reporter constructs and observed that RPRD2 depletion increased transcription of constructs both with and without an intron. We next investigated transcription from integrated proviruses and found no effect of RPRD2 depletion using several different systems. Lastly, we measured transcription of endogenous retroelements and found that RPRD2 depletion did not affect transcription of LINE-1 or HERV-K. Finally, we investigated whether RPRD2 regulates production of IFN in response to nucleic acid species or affects transcription of IFN-stimulated genes. We found that depletion of RPRD2 had no effect on IFN production or ISG expression. Together, our findings demonstrate how regulation of transcription is not universal for host genes, integrated provirus, unintegrated plasmid and endogenous retroviruses, and confirmed that although RPRD2 governs cellular transcription, it does not regulate transcription of HIV-1 provirus or of endogenous retroelements.

## INTRODUCTION

The latent reservoir remains the critical barrier to HIV cure (reviewed in (1)). Despite the effectiveness of antiretroviral therapies, cessation of treatment leads to rapid viral rebound (2,3). However, even with undetectable viral load, persistent transcription of integrated proviruses leads to production of viral transcripts and proteins which contribute to inflammation and co-morbidities for people with HIV (4). A vital question in the field is how and to what extent transcription of viral transcripts is differentially regulated to host transcripts.

RPRD2 (Regulation of nuclear Pre-mRNA Domain containing 2, also known as REAF, reviewed in (5)) has been characterized as a generalized lentiviral restriction factor (6) via inhibition of reverse transcription (6–9), limiting the completion of proviral DNA synthesis and integration of the viral genome (6,8,9). The proposed mechanism of restriction was RPRD2 interaction with DNA:RNA products of reverse transcription as viral both RU5 and late reverse transcripts were enriched in an RPRD2 immunoprecipitation (6). Interestingly, RPRD2 was also precipitated with poly(A)-specific oligo(dT), but treatment with DNase prevented this precipitation, suggesting that RPRD2 specifically interacts with complexes containing both RNA and DNA (6).

However, RPRD2 has also been characterized as a negative regulator of global transcription (10) and interacts with RNA polymerase II (10–12), R-Loops (13), as well as being identified as an RNA-binding protein in several studies (14–19). Furthermore, RPRD2 interacts with members of the HUman Silencing Hub (HUSH) complex (20) which silences transcription of endogenous retroelements and integrated retroviruses (21–26) and the Polymerase-Associated Factor 1 (PAF1) complex (27–29), a multi-functional regulator of both host and viral transcription (21,30–34). Interestingly, members of both the HUSH and PAF1 complexes were identified in the screen for HIV-1 restriction factors that first identified RPRD2 (21). Finally, DNA:RNA hybrids are produced during reverse transcription (35), and also DNA repair (36) and transcription (37). Given this evidence, we reasoned that in addition to inhibiting reverse transcriptase, RPRD2 may regulate nascent transcription of the HIV provirus, potentially via DNA:RNA binding.

Similar to retroviruses, endogenous retroelements encode their own reverse transcriptases and are often transcribed by RNA polymerase II (38). Endogenous retroelements comprise over 40% of the human genome (39,40) and include human endogenous retroviruses (HERVs), the remnants of ancient retroviral infections (41) and long interspersed nuclear elements (LINEs), the most abundant retrotransposons in the human genome (40,42). HERVs, like HIV provirus, contain Long Terminal Repeat (LTR) promoters that can recruit host-cell machinery to drive transcription but are tightly regulated by the cell to suppress their function (43–45). LINEs do not possess LTRs but can still recruit RNA polymerase II and initiate transcription (46). Both HERVs and LINEs are expressed in human cells in a tissue- and developmental stage-specific manner (43,47–49). Given the potential for endogenous retroelement expression to cause genetic disruption, or be immunogenic and influence disease pathology, their transcription is suppressed via several host defense pathways (40,44,45), for example the HUSH complex suppresses LINE-1 transcription (22,26). In a genome-wide screen to identify genes involved in the control of LINE-1 retrotransposition, RPRD2 was identified as the fourth most suppressive hit, following the HUSH complex member TASOR which was the third (25). We therefore also wanted to investigate the role of RPRD2 in regulating transcription of retrotransposons including LINEs and HERVs.

Here, we performed a systematic investigation into RPRD2-mediated regulation of HIV-1 and endogenous retroelement transcription. Using a combination of experimental and computational methods we identified putative RNA binding domains within RPRD2 and confirmed binding to DNA:RNA hybrids. We identified that RPRD2 interacts with the transcription regulator PAF1 and the HUSH complex member TASOR independently, suggesting distinct roles, and GFP-RPRD2 localizes to nuclear speckles, placing RPRD2 at the site of cellular and viral transcription. We investigated the effect of RPRD2 on transcription in several contexts and found that RPRD2 negatively regulates transcription of plasmid-borne HIV-1 LTR-driven reporter constructs but had no impact on transcription of integrated proviruses or endogenous retroelements. Finally, we investigated the role of RPRD2 in the interferon (IFN) response to nucleic acid and found that depletion of RPRD2 had no effect on IFN production or transcription of IFN-stimulated genes. Together, our findings demonstrate that regulation of transcription varies between host genes, integrated provirus, unintegrated plasmid and endogenous retroviruses, and confirmed that although RPRD2 governs cellular transcription, does not regulate HIV-1 or retroelement transcription.

## RESULTS

### RPRD2 binding of DNA:RNA hybrids and dsRNA

RPRD2 has been identified in several screens for RNA binding proteins (14–19) in addition to evidence that it binds DNA:RNA hybrids (6,13) but the interaction interface between the protein and nucleic acids remains unknown. To identify regions of RPRD2 most likely to participate in RNA recognition, we integrated evidence from previously published experimental studies with computational prediction methods and structural interface analyses. RNA-binding domain mapping (RBDmap) (50,51) combined oligo(dT) RNA capture with UV crosslinking and mass spectrometry, to identify RNA-binding peptides, thereby mapping RNA binding regions within proteins. RBDmap highlighted two RNA binding sites within the C-terminal region of RPRD2 (**Figure 1A, top panel**). Computational prediction using the hybrid ensemble RBP classifier (HydRA) model (52) identified the N-terminal CTD-interacting domain (CID) as the principal contributor to RNA-binding activity (**Figure 1A, middle panel**), whereas our recently developed RNA binding protein language model (RBP-LM) (manuscript in preparation) predicted RNA-binding regions spanning the CID and CREPT domains together with additional intrinsically disordered segments located within the C-terminus (**Figure 1A, bottom panel**).

**Figure 1.**
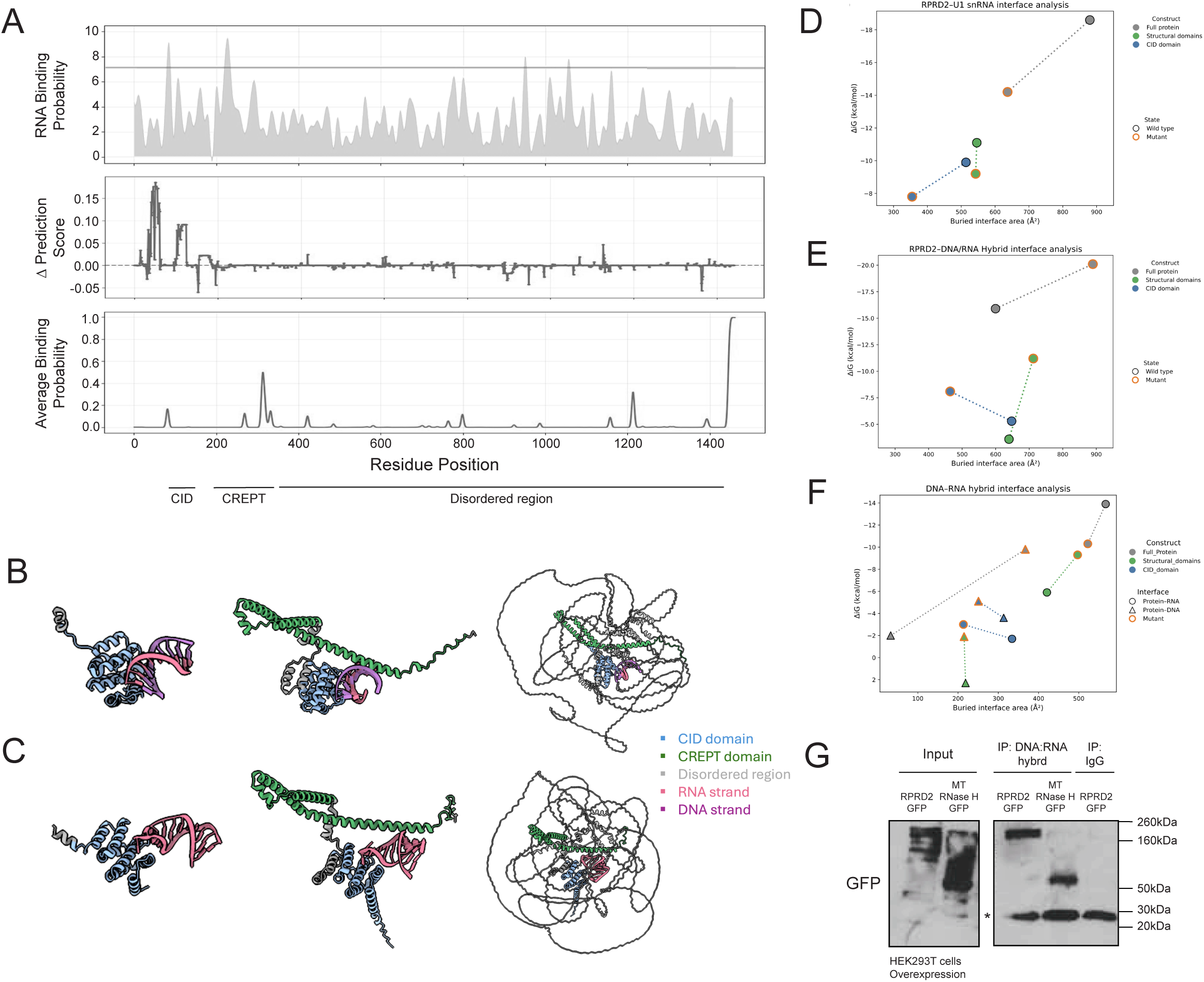
RPRD2 binds DNA:RNA hybrids. **A)** RNA-binding probability score along length of RPRD2 protein predicted experimentally by RBDmap (50) (top panel), or computationally using HydRA (52) (middle panel), or our novel RBP-LM model (manuscript in preparation) (bottom panel). Plots show (top) RNA-binding probability (RNA-bound / released [log2-ratio]), (middle) Δ Prediction score (Occluded-Original), and (bottom) Average Binding Probability on the y axes respectively, and amino acid position of RPRD2 on the x axis. The bold line on the RBDmap plot indicates likelihood cut off. Previously characterized domains of RPRD2 are shown below; CID = RNA polymerase II carboxy-terminal domain (CTD)-Interacting Domain, CREPT = cell-cycle alteration and expression-elevated protein in tumor family domain. **B)** Complex structure of RPRD2 CID domain (left panel; blue), CIDD and CREPT domains (middle panel; blue and green) and full RPRD2 protein (right panel) with DNA:RNA hybrid molecule or **C)** with U1 snRNA stem-loop structure. RNA is colored pink while DNA is colored purple in DNA/RNA hybrid. **D)** Interface energetics of wild-type and mutant RPRD2 nucleic acid complexes. Relationship between buried interface area and predicted binding free energy (ΔiG) for wild-type and mutant RPRD2 models interacting with U1 snRNA, **E)** DNA/RNA hybrid, or **F)** isolated DNA-RNA hybrid interfaces. Data are shown for full-length protein (grey), structured domains (green), and CID domain (blue) constructs. Lower ΔiG values indicate more favorable predicted binding energetics, while larger buried surface areas represent greater interfacial contacts. Lines connect corresponding wildtype (grey outline) and mutant (orange outline) models within each construct. **G)** Protein immunoprecipitation of DNA:RNA hybrids. HEK293T cells were transfected with either GFP-RPRD2 or GFP-RNaseH MT as positive control. Protein was isolated 48 h later and incubated with S9.6 antibody specific for DNA:RNA hybrids. Input total lysate and immunoprecipitate were resolved by SDS-PAGE and probed for GFP expression using immunoblot. Star (*) indicates band of antibody used in immunoprecipitation.

To further evaluate these predictions from a structural perspective, protein-RNA complexes generated using AlphaFold (53) were analyzed using UCSF ChimeraX (54). Three RPRD2 structures were analyzed; the full-length RPRD2 protein, a construct encompassing the CREPT and CID domains, and an experimentally resolved structure of CID only (11). RPRD2 has been suggested to regulate transcription of U1 small nuclear RNA (snRNA) (10), so a dsRNA stem loop from U1 was modelled in addition to a DNA:RNA hybrid. Intermolecular contacts between RPRD2 and a DNA:RNA hybrid (**Figure 1B**) or the U1 snRNA stem-loop 4 (**Figure 1C**) were identified using a 4 Å distance threshold, allowing residues located at the predicted interaction interfaces to be systematically catalogued (**Table S2**). Contact analyses performed across all 3 protein constructs consistently revealed clusters of interacting residues within the N-terminal CID and CREPT domains, in agreement with the regions predicted by HydRA, RBP-LM, and RBDmap experimental evidence.

To further evaluate the functional relevance of the predicted RNA-binding regions, targeted mutational analysis was performed on key interface residues identified as potential binding hotspots. Three positively charged residues frequently identified at the interaction interfaces were selected: Arg90, Arg130, and Lys304. Each residue was individually substituted with glutamic acid (R90E, R130E, and K304E) to reverse the local electrostatic charge while maintaining minimal changes to residue size. Structural confidence of all mutant proteins was compared with their respective wild-type counterparts using residue-wise pLDDT scores (**Figure S1A-C**) indicating that the mutations did not substantially alter the predicted protein architecture and any subsequent differences in nucleic acid interaction are more likely to arise from disruption of intermolecular contacts than from global structural destabilization. PAE values (**Figure S1D**) to evaluate the confidence of the predicted intermolecular interfaces and mean ipTM scores (**Figure S1E**) to assess changes in interface confidence, remained largely comparable between wild-type and mutant complexes for both U1 snRNA and the DNA:RNA hybrid, indicating that AlphaFold retained similar confidence in the overall placement of the interacting molecules following mutation. The effect of these mutations on RPRD2-RNA interaction was subsequently analyzed using the PDBePISA server (55) to characterize intermolecular interfaces and estimate interaction stability. Analysis of U1 snRNA binding demonstrated a consistent reduction in interface stability following mutation across all three RPRD2 constructs (**Figure 1D**). Compared with their corresponding wild-type complexes, mutant proteins formed smaller buried interfaces that were accompanied by less favorable interface free energies. The full-length protein exhibited the largest decrease in both buried interface area and binding energy, while similar trends were observed for the structured-domain and isolated CID constructs. Together, these results indicate that substitution of Arg90, Arg130 and Lys304 weakens the predicted interaction between RPRD2 and U1 snRNA, supporting the functional importance of these positively charged residues in RNA recognition.

Conversely, analysis of DNA:RNA hybrid binding showed that rather than reducing interface stability; mutant complexes generally displayed larger buried interface areas and more favorable ΔiG values than their wild-type counterparts (**Figure 1E**). These observations suggest that disruption of the selected residues does not impair overall recognition of the hybrid substrate and may instead promote alternative intermolecular contacts. To distinguish interactions with the individual nucleic acid strands, DNA and RNA interfaces within the hybrid complexes were analyzed separately (**Figure 1F**). This analysis revealed that mutations consistently reduced the stability of the protein-RNA interface, as reflected by decreased buried surface areas and less favorable binding energies across all three protein constructs. In contrast, interactions with the DNA strand were generally strengthened following mutation, producing larger interface areas and more favorable ΔiG values. These findings indicate that the selected hotspot residues contribute preferentially to RNA recognition, whereas interaction with the DNA component of the hybrid can be maintained or compensated through alternative contacts. Therefore, these results support the hypothesis that the CID and CREPT regions participate in RNA binding while also contributing to the broader nucleic acid recognition properties of RPRD2.

Finally, we confirmed experimentally that RPRD2 binds DNA:RNA hybrids naturally occurring in uninfected cells. HEK293T cells were transfected with either GFP-RPRD2 or GFP-RNase H with a mutated cleavage site as a positive control. After 48 hours, protein was isolated, and DNA:RNA hybrids were immunoprecipitated using the specific but largely sequence-independent antibody S9.6 (56). Bound protein was eluted and resolved by SDS-PAGE. Immunoblotting for GFP revealed that both the mutant RNase H and RPRD2 were present in the immunoprecipitated fraction (**Figure 1G**), indicating they were bound to the DNA:RNA hybrids.

Together these results provide experimental and computational evidence for the binding of RPRD2 to both RNA and DNA:RNA hybrids and propose the likely interaction domains within the N terminal of the protein. It is highly likely that the intrinsically disordered C terminal of RPRD2 also plays a role in binding nucleic acids since this is the case for many RNA-binding proteins (57).

### RPRD2 interacts with transcriptional regulators and localizes to nuclear speckles

RPRD2 was first characterized as an RNA polymerase II binding protein, interacting with its C-terminal tail (11). The Polymerase-Associated Factor 1 (PAF1) complex also binds to the RNA polymerase II C tail and has essential cellular roles including both transcription activation and repression, and regulation of transcription elongation, and R-loop formation (28,33). The eponymous protein PAF1 has also been characterized as an HIV restriction factor (21,58) and to mediate antiviral gene expression in response to Influenza A infection (59). RPRD2 has previously been identified as an interaction partner of PAF1 complex members LEO1 and CTR9 (27), and PAF1 itself (28,29). We were therefore interested in whether RPRD2 interacts with PAF1. To investigate, we performed immunoprecipitation on Jurkat cell lysates of either endogenous RPRD2 or endogenous PAF1 to isolate interacting proteins and found that RPRD2 was present in the PAF1 immunoprecipitate (**Figure 2A**), suggesting they interact either directly or indirectly. RPRD2 has also been found to interact with and even suggested to be a member of the HUman Silencing Hub (HUSH) complex (20), which inhibits transcription of DNA such as retroelements (25,26), endogenous retroviruses (60,61), and integrated HIV-1 provirus (23,62–64). We therefore repeated the immunoprecipitation and additionally included immunoprecipitation of TASOR (**Figure 2B**). We found that TASOR was co-immunoprecipitated with RPRD2 but not with PAF1, suggesting RPRD2 interacts with each of these complexes independently. This is in agreement with protein interaction predictions which do not show evidence of PAF1-TASOR interactions despite several other interactions between other members of both complexes (**Figure 2C**), highlighting the close association of RPRD2, RNA polymerase II and the PAF1 and HUSH complexes.

**Figure 2.**
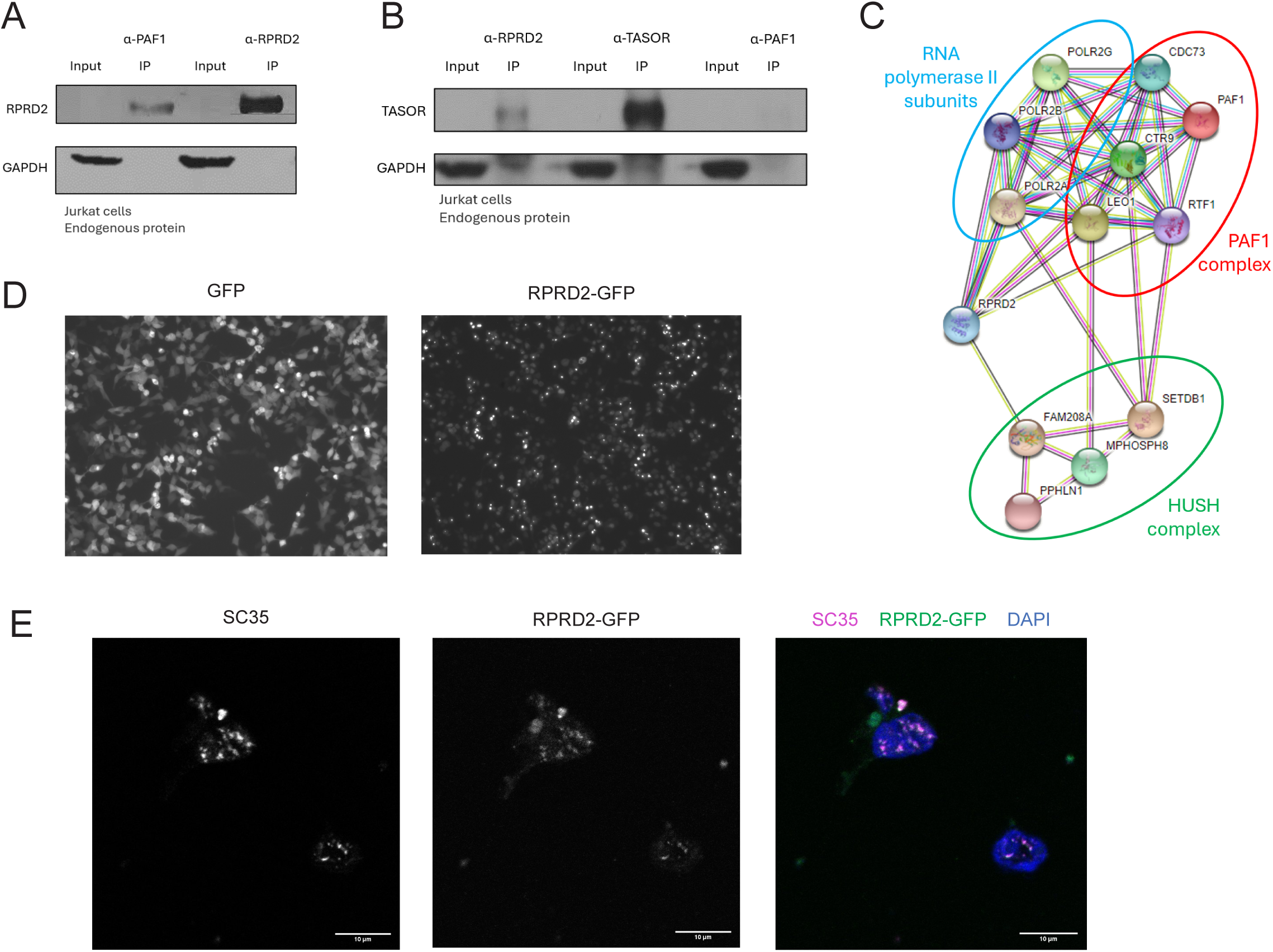
RPRD2 interacts with transcription regulators and localizes to nuclear speckles. **A)** Immunoprecipitation of endogenous PAF1 or RPRD2. Jurkat cells were harvested and protein lysate incubated with PAF1 or RPRD2 antibody, followed by immunoblot and probe for RPRD2 in input and IP samples. GAPDH was probed as loading control. **B)** Immunoprecipitation of endogenous TASOR, PAF1 or RPRD2. Jurkat cells were harvested and protein lysate incubated with TASOR, PAF1 or RPRD2 antibody, followed by immunoblot and probe for TASOR in input and IP samples. GAPDH was probed as a loading control. **C)** Experimental and predicted interactions between RPRD2, HUSH complex members, PAF1 complex members and RNA polymerase II. Protein interaction network was generated using the STRING database (https://string-db.org/). Line color indicates type of connection between proteins; cyan = known interaction from curated databases, magenta = experimentally determined interaction, black = genes that are co-expressed, yellow = proteins found together in literature by text mining, and green = predicted interaction based on gene neighborhood. **D)** Immunofluorescence imaging of GFP-RPRD2 overexpression and GFP control. HEK293T cells were transfected with GFP-RPRD2 plasmid or GFP only vector control. Two days later, cells were fixed and imaged using confocal microscopy. **E)** Immunofluorescence imaging of GFP-RPRD2 overexpression and SC35 nuclear speckle marker. HEK293T cells were transfected with GFP-RPRD2 plasmid, and two days later, cells were permeabilized, incubated with SC35 specific antibody, stained with DAPI and then fixed and imaged using confocal microscopy.

Transcription takes place in the nucleus inside nuclear speckles; liquid phase separated foci with a concentration of transcription factors and machinery. HIV-1 replication complexes also accumulate in nuclear speckles and integrate into speckle-associated genomic domains (65). The formation and maintenance of this phase separated speckles is regulated by interactions between disordered domains of several proteins (66–68). RPRD2, like many RNA-binding proteins (57), is highly disordered. We therefore were next interested to see if RPRD2 is present in these sites of transcription. We transfected HEK293T cells with GFP-RPRD2 or a GFP only vector control and used immunofluorescence to visualize the location of each protein (**Figure 2D**). GFP alone was dispersed throughout the cell, whereas GFP-RPRD2 was mostly localized to the nucleus and showed bright foci within the nuclei. We next imaged the nuclear speckle marker SC35 (SRSF2) (69,70) in addition to GFP-RPRD2 and found that the GFP-RPRD2 foci co-localized with the SC35 foci (**Figure 2E**).

These findings suggest RPRD2 localizes to SC35-enriched nuclear foci, suggesting that RPRD2 is found within nuclear speckles where host and viral transcription take place. These results, in addition to previous work, place RPRD2 at the site of transcription of both host and viral RNA and implies RPRD2 is an interactor of both the PAF1 and HUSH complexes, well characterized regulators of viral and endogenous retroelement transcription respectively. Further, these results suggest that RPRD2 binds TASOR and the PAF1 complex independently, perhaps suggesting two distinct roles in the vicinity of transcription, or targeting of distinct transcriptional contexts.

### RPRD2 knockdown increases HIV LTR-driven transcription from reporter plasmids

We next investigated the role of RPRD2 on HIV transcription. To decouple transcription regulation from any effects of RPRD2 on reverse transcription, we used an LTR-driven GFP reporter construct expressed form a plasmid (pHIV-1 LTR-Tat-IRES-GFP) (71). We manipulated protein levels of RPRD2 or TASOR as a control in HEK293T cells, using siRNA to knockdown RPRD2, TASOR or non-targeting control and plasmid transfection to overexpress RPRD2, TASOR or empty vector control. Quantification of mRNA levels by qRT-PCR also showed the appropriate decreased or increased levels, respectively (**Figure 3A**). Immunoblotting confirmed a decrease of RPRD2 or TASOR protein levels respectively, or both in double knockdown cells, and an increase in RPRD2 levels in overexpression cells (**Figure 3B**). Although we observed a large increase in TASOR mRNA in the overexpression sample, we did not observe an increase in TASOR protein levels. Interestingly, we also noted that RPRD2 overexpression led to a decrease in TASOR protein level, possibly suggesting a feedback mechanism. Each condition was co-transfected with the GFP expressing construct driven by an HIV-1 LTR promoter that was used originally to create J-Lat cell lines (71). The percentage of GFP positive cells was measured 48 h later by flow cytometry (**Figure 3C**). Knockdown of RPRD2 led to an increase in the percentage of GFP positive cells, suggesting RPRD2 has an inhibitory role in regulating LTR-driven transcription from plasmid DNA. Knockdown of TASOR led to a larger increase, whilst double knockdown led to a similar increase as TASOR alone, suggesting the effect may be due to perturbation of the same pathway. In contrast, overexpression of either RPRD2 or TASOR did not significantly alter the percentage of GFP positive cells.

**Figure 3.**
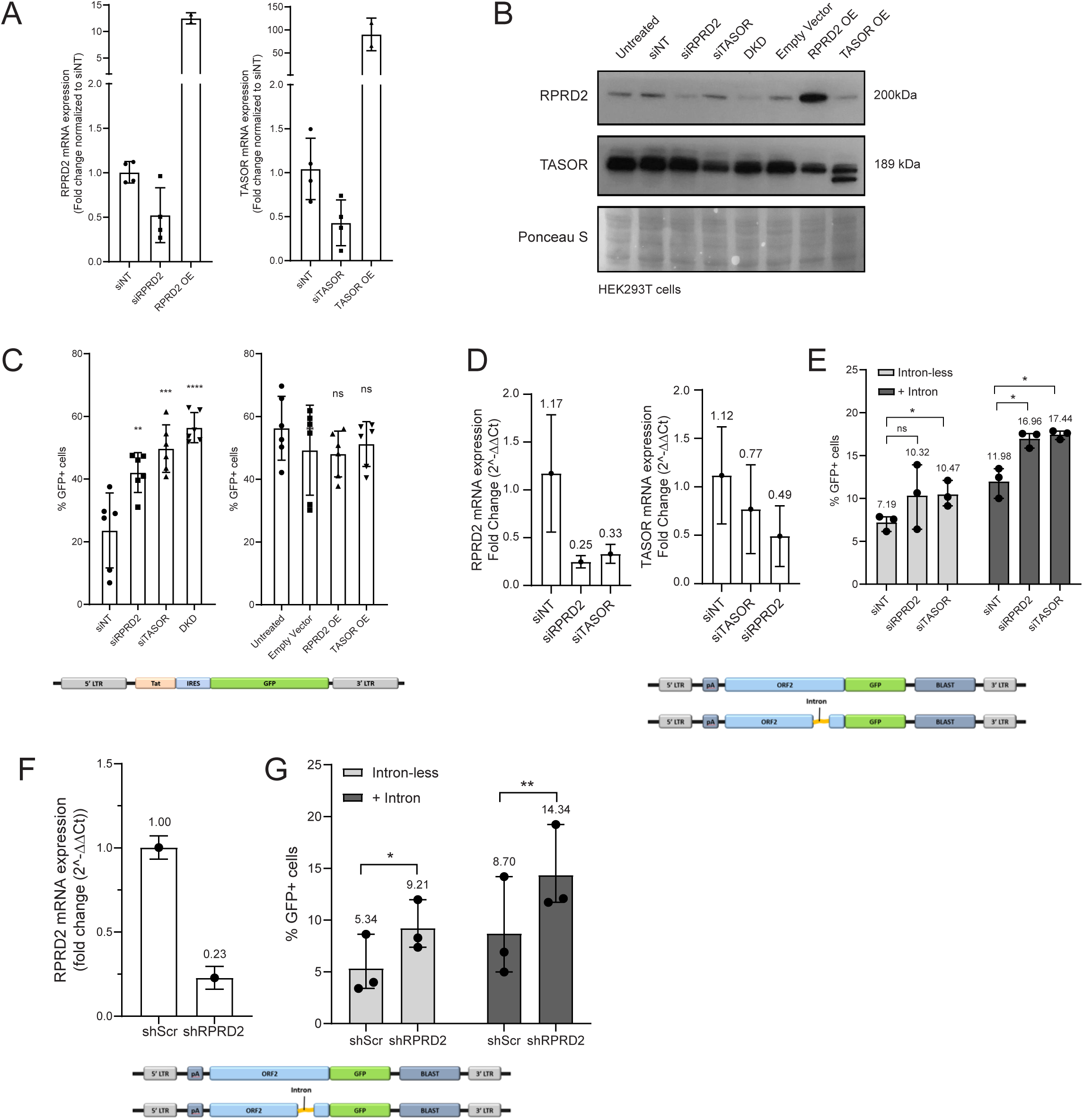
RPRD2 knockdown increases HIV LTR-driven transcription from plasmid. **A)** Knockdown or overexpression efficiency of RPRD2 and/or TASOR mRNA. HEK293T cells were transfected with either siRNA targeting RPRD2, TASOR, both or non-targeting control, or overexpression vectors for RPRD2 or TASOR. After 48 h, RNA was isolated and TASOR transcript levels were quantified by qRT-PCR relative to the NT control. **B)** Knockdown or overexpression efficiency of RPRD2 and/or TASOR proteins. HEK293T cells were treated as in A, protein was isolated, run on immunoblot and probed with RPRD2 or TASOR antibodies. Ponceau S stain is shown as a loading control. **C)** Effect of RPRD2 and TASOR knockdown or overexpression on HIV LTR-driven transcription of reporter construct. HEK293T cells were treated as in A, and co-transfected with the pLTR-Tat-IRES-GFP reporter plasmid. Transcription was measured 48 h later as percent GFP positive cells by flow cytometry. **D)** Confirmation of knockdown of RPRD2 and TASOR mRNA by siRNA. HEK293T cells were transfected with siRNA targeting RPRD2, TASOR, or non-targeting control. RNA was isolated 48 h after transfection, and RPRD2 and TASOR transcript levels were quantified by qRT-PCR relative to the NT control. **E)** Effect of RPRD2 and TASOR knockdown on HIV LTR-driven transcription of intron-containing or intron-less reporter. HEK293T cells were treated as in D, and co-transfected with either the intron-less or intron-containing HIV-1 LTR-driven ORF2 GFP reporter plasmid. Transcription was measured 48 h later as percent GFP positive cells by flow cytometry. **F)** Knockdown efficiency of stably integrated, inducible RPRD2 shRNA. HeLa cells that harbor an IPTG-inducible shRNA targeting either RPRD2 (i-shRPRD2-HeLa) or scramble (scr) control (i-shScr-HeLa) were treated with IPTG for 72 h. mRNA was isolated and RPRD2 transcript levels quantified by qRT-PCR. **G)** Effect of RPRD2 knockdown on HIV LTR-driven transcription of intron-containing or intron-less reporter plasmid. IPTG-treated i-shRPRD2-HeLa or i-shScr-HeLa cells were transfected with either the intron-less or intron-containing HIV-1 LTR-driven ORF2 GFP reporter plasmid. Transcription was measured 48 h later as percent GFP positive cells by flow cytometry.

Since the HUSH complex preferentially binds and silences intron-less genetic elements (26,72), we next wanted to characterize whether RPRD2 is also able to regulate transcription of these DNA species. We therefore repeated the siRNA knockdown of RPRD2 or TASOR in HEK293T cells followed by transfection of lentiviral HIV-1-derived intron-less or intron-containing reporter plasmids. Quantification of mRNA levels by qRT-PCR showed decreased RPRD2 transcripts in siRPRD2 cells, and also a decrease in siTASOR cells (**Figure 3D**). TASOR transcripts were decreased in siTASOR cells, and also in siRPRD2 cells, perhaps suggesting a feedback mechanism between transcription or RNA stability of the two genes. Expression of the intron-containing reporter was higher than the intron-less construct regardless of siRNA treatment (**Figure 3E**). Depletion of TASOR led to an increase in the percentage of GFP positive cells compared to non-targeting for both the intron-less and intron-containing constructs. Knockdown of RPRD2 led to a similar increase in the percentage of GFP positive cells for the intron-containing construct but did not significantly impact the intron-less construct. We also transfected these constructs into HeLa cells harboring either an inducible shRNA targeting scramble control (i-shScr-HeLa) or RPRD2 (i-shRPRD2-HeLa) (previously described (8,9), 24 h after isopropyl-β-d-1-thiogalactopyranoside (IPTG) treatment to induce knockdown. Validation of RPRD2 knockdown by qRT-PCR and flow cytometry analysis of GFP reporter expression were carried out 48 hours later. Decrease of RPRD2 mRNA was significant (**Figure 3F**). As seen previously, the expression of the intron-containing reporter was higher than the intron-less construct in both the i-shScr-HeLa and i-shRPRD2-HeLa cells (**Figure 3G**). We observed a significant increase in the percentage of GFP cells in RPRD2 depleted cells compared to i-shScr-HeLa cells for both the intron-less and the intron-containing constructs.

Together, these findings suggest that RPRD2 is negatively regulating LTR-driven transcription from transfected plasmids. The lack of additive effect when both RPRD2 and TASOR were depleted suggests RPRD2 may be working via the same pathways as TASOR, potentially as part of the HUSH complex. In addition, the loss of both RPRD2 and TASOR mRNA when either was depleted with siRNA suggests some feedback mechanism between transcription of the two genes.

### RPRD2 does not regulate integrated HIV LTR-driven transcription

Thus far, LTR-driven transcription measured has been from reporter plasmid constructs only. Whilst transcription can occur from unintegrated viral DNA (73–75), the majority of transcription originates from integrated provirus. We therefore wanted to measure the effect of RPRD2 on integrated provirus transcription. We created inducible shRNA cell lines from two different J-Lat clones with HIV-1 proviral integration in different loci, using the same shRNAs targeting RPRD2 or non-targeting control as in the i-shRPRD2-HeLa and i-shScr-HeLa cells. After 72 h IPTG treatment, we confirmed depletion of RPRD2 at the mRNA level compared to non-targeting control using qRT-PCR (**Figure 4A**). Cells were then stimulated using TNFα to activate the integrated HIV-1 reporter, and 24 h later, the percentage of GFP positive cells were measured by flow cytometry as a proxy for transcription of the integrated provirus. We found no change to the percentage of GFP positive cells for either J-Lat clone in the RPRD2 depleted cells compared to the control cells (**Figure 4B**), suggesting RPRD2 does not regulate transcription of integrated provirus in these cells.

**Figure 4.**
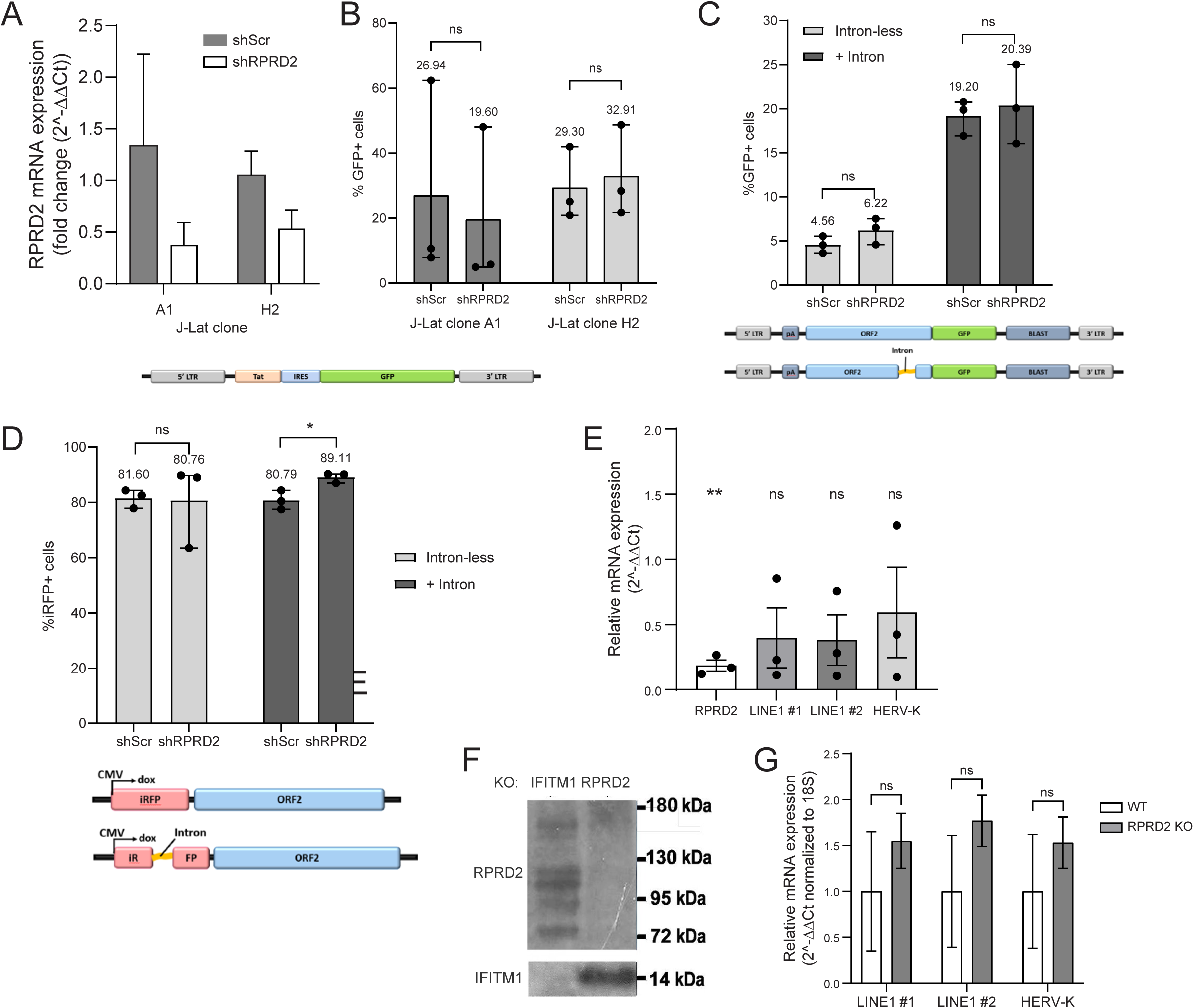
RPRD2 does not regulate integrated HIV LTR-driven transcription or endogenous retroelement transcription. **A)** Knockdown efficiency of stably integrated, inducible RPRD2 shRNA in J-Lats. J-Lat clones A1 and H2 cells were transduced with the shRNA cassettes targeting RPRD2 or scramble (scr) control, and cells with successful integration selected by puromycin treatment. Inducible shRNA J-Lat cells were then treated with IPTG for 72 h, mRNA isolated, and RPRD2 mRNA levels quantified by qRT-PCR. **B)** Effect of RPRD2 knockdown on HIV LTR-driven transcription of integrated provirus. J-Lat cells described in A that harbor an IPTG-inducible shRNA targeting either RPRD2 or shScr control were treated with IPTG for 72 h. After 48 h, cells were additionally stimulated with TNFα to activate latent provirus. Transcription was measured as percent GFP positive cells by flow cytometry. **C)** Effect of RPRD2 knockdown on HIV LTR-driven transcription of integrated intron-containing or intron-less reporter. i-shRPRD2-HeLa or i-shScr-HeLa cells were treated with IPTG for 72 h. After 48 h, cells were transduced with either the intron-less or intron-containing HIV-1 LTR-driven ORF2 GFP reporter plasmid. Transcription was measured as percent GFP positive cells by flow cytometry. **D)** Effect of RPRD2 knockdown on transcription of integrated LINE-1-derived piggyBac intron-containing or intron-less reporter.i-shRPRD2-HeLa or i-shScr-HeLa cells were transduced with a LINE-1-derived piggyBac cassette either with an intron in the RFP reporter or not, and cells with integrated reporter selected by Blasticidin treatment. Cells were then treated with IPTG for 72 h, and transcription was measured as percent RFP positive cells by flow cytometry. **E)** Effect of RPRD2 knockdown on transcription of endogenous LINE-1 and HERV-K. HEK293T cells were depleted of RPRD using shRNA. After 48 h, RNA was extracted and endogenous LINE-1 and HERV-K levels were measured by qRT-PCR compared to siNT control treated cells. **F)** Knockout efficiency of RPRD2 or IFITM1 control following CRISPR-Cas9 editing of K562 cells. **G)** Effect of RPRD2 knockdown on transcription of endogenous LINE-1 and HERV-K. RNA was extracted from RPRD2 knockout and wild type (WT) K562, and endogenous LINE-1 and HERV-K levels were measured by qRT-PCR.

Regulation of proviral transcription is known to be regulated by a complex network of mechanisms, influenced by cellular factors and integration locus (76). We therefore wanted to measure transcription from a wide array of integration loci. We again used the LTR-GFP constructs with and without an intron, but here we transduced them into the i-shRPRD2-HeLa and i-shScr-HeLa cells using a lentiviral vector. Cells were selected with Blasticidin for 7 days, resulting in a mixed population of cells with various reporter integration sites. Cells were then treated with IPTG for 72 hours to induce the shRNA knockdowns and transcription quantified by the percentage of GFP positive cells measured by flow cytometry (**Figure 4C**). Again, we observed that the expression of the intron-containing reporter was higher than the intron-less construct in both i-shScr-HeLa and i-shRPRD2-HeLa cells. There was no difference in the percentage of GFP positive cells in the RPRD2 depleted cells compared to the i-shScr-HeLa for cells transduced with either the intron-less or the intron-containing construct.

Although our previous results indicated that RPRD2 negatively regulates LTR-driven transcription, those experiments utilized plasmid-based reporter constructs. Here we investigated transcription of reporters integrated into the host genome, more physiologically representative of HIV-1 proviral transcription. We saw no changes after RPRD2 depletion regardless of integration site or presence of an intron. These results suggest that RPRD2 does not regulate transcription of integrated viral DNA regardless of intron presence.

### RPRD2 does not regulate transcription of endogenous retroelements

We found no change to integrated provirus transcription in RPRD2 depleted cells. However, RPRD2 has been shown to negatively regulate global transcription (10), and additionally, RPRD2, along with HUSH complex members, were identified as hits in a screen to find regulators of long interspersed nuclear elements (LINEs) retrotransposition in K562 cells (25). We therefore next utilized LINE-1-derived constructs containing an RFP reporter with and without an intron (26). The i-shRPRD2-HeLa and i-shScr-HeLa cells were transfected with each reporter construct plus a piggyBac transposase-expression plasmid to allow integration through the PiggyBac transposon system (26). Cells were selected with Blasticidin for 7 days, followed by IPTG treatment for 72 hours to induce expression of the shRNA, and doxycycline treatment to induce expression of the reporter constructs. After 24 hours, transcription was measured by the percentage of RFP positive cells by flow cytometry (**Figure 4D**). We observed a very small but significant increase in the percentage RFP positive cells in RPRD2 depleted cells with integrated intron-containing reporter, but no change to percentage RFP positive cells in RPRD2 depleted cells with integrated intron-less reporter.

Although RPRD2 depletion did not impact transcription of the integrated intron-less LINE-1 reporter, we wanted to confirm this finding by investigating whether RPRD2 regulates transcription of endogenous LINEs or human endogenous retroviruses (HERVs). We therefore measured endogenous LINE-1 and HERV-K RNA levels by qRT-PCR after depletion of RPRD2 using shRNA in HEK293T cells (**Figure 4E**). Despite efficient knockdown of RPRD2, we saw no significant effect on LINE-1 or HERV-K transcription. Reasoning that this could be a cell type specific mechanism, we next utilized K562 cells, widely used in endogenous retroelement research as they have active HERVs and LINE-1s (77,78), and the cell type used in the LINE-1 screen mentioned above (25). Using CRISPR-Cas9 we targeted RPRD2, or IFITM1 as a control, in K562 cells and confirmed depletion by immunoblot which showed a clear decrease in protein levels of each target (**Figure 4F**). RNA was isolated from WT or RPRD2 knockout cells, and LINE-1 and HERV-K RNA levels were measured by qRT-PCR (**Figure 4G**). Once again, we saw no significant effect on LINE-1 or HERV-K transcription.

Although RPRD2 depletion did lead to a small but significant increase in transcription of the integrated, LINE-1-derived intron-containing reporter, these findings indicate that RPRD2 does not regulate transcription of intron-less endogenous retroelements such as LINEs and HERVs.

### RPRD2 does not impact the interferon response to nucleic acids

Our findings suggest that RPRD2 does not regulate transcription of integrated viral DNA or endogenous retroelements, but RPRD2 is known to interact with RNA polymerase II (11,12,79) and has been found to regulate host transcription (10). We therefore wanted to investigate whether RPRD2 regulation of specific host genes could influence viral infection. Equally we were interested if RPRD2 driven expression of interferon might underlie retroelement control, as multiple ISGs are known to regulate retroelements (80). To investigate the constitutive expression of RPRD2 we accessed RNA and protein expression data from The Human Protein Atlas database of tissue expression (81,82). Both RPRD2 mRNA and protein expression have low cell type specificity, and relative mRNA expression does not correlate to levels of protein expression across tissues (**Figure 5A**). Nevertheless, RPRD2 protein is enriched in immune cells including lymphocytes, as well as ciliated cells, and neurons. To investigate immune cells more closely, we interrogated the Monaco dataset (83) which consists of RNA-seq profiling of 29 immune cell types. Among immune cells, RPRD2 was found to be most highly expressed in T cells, followed by B cells and NK cells (**Figure 5B**).

**Figure 5.**
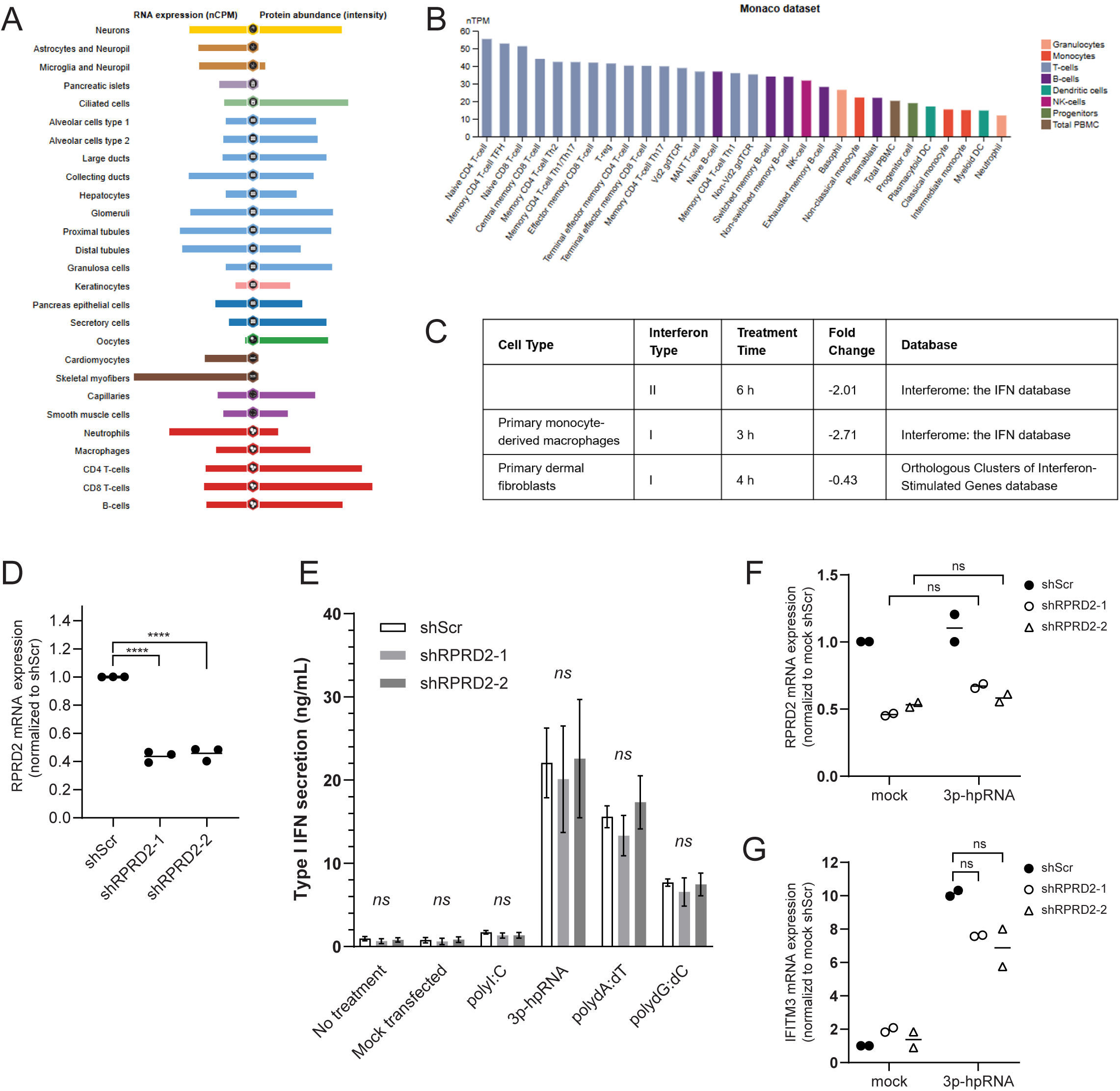
RPRD2 does not regulate interferon type I production. **A)** Expression level of RPRD2 RNA and protein across cell types. Data were retrieved from The Human Protein Atlas database of tissue expression (81,82). **B)** Expression level of RPRD2 RNA across immune cell subtypes. Data were retrieved from The Human Protein Atlas database of tissue expression. **C)** Effect of interferon (IFN) treatment on RPRD2 RNA levels. RPRD2 RNA expression fold change was retrieved from three studies that treated various cell types with type I and/or type II IFN. Data were retrieved from Interferome: the IFN database (85) and Orthologous Clusters of Interferon-Stimulated Genes database (86). **D)** Knockdown efficiency of stably integrated, inducible RPRD2 shRNA in THP-1 cells (i-shScr-THP-1, i-RPRD2-1-THP-1 and i-RPRD2-2-THP-1). Cells were transduced with the shRNA cassettes targeting RPRD2 or scramble (shScr) control, and cells with successful integration selected by Blasticidin treatment. Inducible THP-1 cells were then treated with IPTG for 72 h, mRNA isolated, and RPRD2 transcript levels quantified by qRT-PCR. **E)** Effect of RPRD2 on IFN type I production in response to nucleic acid agonists. The inducible shRNA THP-1 cells were subsequently transfected with various nucleic acid IFN agonists; polyI:C, 3p-hpRNA, polydA:dT and polydG:dC, or transfection reagent alone or untreated. After 24 hours, supernatant was collected from the cells and the type I IFN secretion levels quantified using the HEK Blue reporter system (Invivogen). **F)** Effect of 3p-hpRNA treatment and RPRD2 depletion on RPRD2 and **G)** IFITM3 mRNA levels. The i-shScr-THP-1, i-RPRD2-1-THP-1 and i-RPRD2-2-THP-1 cells were induced with IPTG followed by treatment with 3p-hpRNA or transfection reagent alone. After 24 hours, RNA was isolated and IFITM3 and RPRD2 mRNA levels quantified by qRT-PCR.

Although RPRD2 is a generalized retroviral restriction factor (6), it is not an interferon (IFN)-stimulated gene (ISG). In prior work, treatment of THP-1 cells or monocyte derived macrophages with IFN α, β or γ did not increase RPRD2 mRNA or protein levels (9) demonstrating RPRD2 is not IFN-inducible. We accessed the Immune Dictionary, a compendium of single-cell transcriptomic profiles of 17 immune cell types in response to each of 86 cytokines in mouse lymph nodes in vivo (84) to investigate whether RPRD2 expression was influenced by any other signaling molecules. Despite over 1,400 cytokine–cell type combinations, RPRD2 was not identified in any of the datasets. We also searched for RPRD2 in the Interferome: the IFN database (85), which collates datasets from types I, II and III IFN-treated cells, mice or humans, and the Orthologous Clusters of Interferon-Stimulated Genes database (86). RPRD2 mRNA expression was found to be significantly decreased in response to both type I and type II IFN in both databases (**Figure 5C**).

The ability of RPRD2 to bind nucleic acids suggests it could function as an innate immune sensor. The findings that expression of RPRD2 was diminished within 3-6 hours of IFN stimulation is consistent with this function, as IFN signaling frequently induces feedback mechanisms that attenuate nucleic acid sensing pathways once an antiviral response has been initiated (6). We therefore wanted to investigate whether RPRD2 affects the IFN response to nucleic acids. We used inducible shRNA clones of THP-1 cells with the scramble control shRNA (i-shScr-THP-1 cells) or 2 independent shRNAs targeting RPRD2 (i-RPRD2-1-THP-1 and i-RPRD2-2-THP-1 cells) as described above. After 72 h IPTG treatment, RNA was isolated and depletion of RPRD2 at the mRNA level compared to i-shScr-THP-1 control cells was confirmed using qRT-PCR (**Figure 5D**). We then measured the type I IFN secretion level in response to several synthetic nucleic acid agonists after RPRD2 depletion. After 72 h IPTG treatment, cells were transfected with double-stranded RNA polyI:C (TLR3, RIG-I/MDA5 and PKR agonist), single-stranded RNA 3p-hpRNA (RIG-I agonist), and double-stranded DNAs polydA:dT (AIM2 and DAI agonist), and polydG:dC (cGAS and DAI agonist), treated with transfection reagent alone or untreated. After 24 hours, medium was collected from the cells, and type I IFN level quantified by HEK Blue reporter system (**Figure 5E**). All treatments led to an increase in IFN secretion, although this varied between treatments with 3p-hpRNA eliciting the largest response, and polyI:C treatment resulting in a minimal response. Knockdown of RPRD2 by either shRNA had no significant effect on the level of IFN secretion to each nucleic acid species. We repeated this with varying concentrations of 3p-hpRNA (**Figure S2A**) and polyI:C (**Figure S2B**), and with both low and high molecular weight polyI:C species (**Figure S2C**) and observed no difference in RPRD2 knockdown cells compared to scramble control, suggesting that RPRD2 does not act as innate immune sensor of nucleic acid nor regulate the IFN production pathway.

Although RPRD2 did not affect IFN production, downstream IFN signaling triggers the transcription of hundreds of ISGs which restrict viral replication by various mechanisms (87). To determine if RPRD2 regulates transcription of downstream ISGs, we measured expression of the IFNα-induced HIV-1 restriction factor IFITM3 in cells depleted of RPRD2. The i-shScr-THP-1, i-RPRD2-1-THP-1 and i-RPRD2-2-THP-1 cells described above were induced with IPTG followed by treatment with 3p-hpRNA which produced the strongest IFN response of agonists tested (**Figure 5E**). After 24 hours, RNA was isolated and IFITM3 and RPRD2 mRNA levels quantified by qRT-PCR. The i-RPRD2-2-THP-1 cells showed decreased levels of RPRD2 mRNA; however the level of RPRD2 mRNA was not affected by 3p-hpRNA treatment in either control or knockdown cells (**Figure 5F**). As expected, we observed a strong increase in IFITM3 mRNA upon 3p-hpRNA treatment (**Figure 5G**). However, depletion of RPRD2 did not significantly affect IFITM3 expression, in either 3p-hpRNA treated or control cells.

Although RPRD2 is highly expressed in immune cells, and is downregulated rapidly following infection and IFN stimulation, our findings indicate that it does not act as a nucleic acid sensor, regulate IFN production or govern transcription of the representative ISG IFITM3.

## DISCUSSION

Transcriptional regulation is highly dynamic, tightly controlled, and incredibly complex. Understanding the mechanisms that govern transcription in different contexts remains poorly understood. For example, comprehending the regulation of transcription at the HIV-1 provirus is crucial to overcome the persistent viral reservoirs that require lifelong treatment and contribute to co-morbidities even in individuals on suppressive antiretroviral therapy. In this study, we investigated the role of the HIV-1 restriction factor RPRD2 in repressing the transcription of RNA including HIV-1 provirus, endogenous retroviruses, and transposable elements. RPRD2 has previously been characterized as an inhibitor of reverse transcription (6) and suggested to bind various nucleic acid species including poly(A) RNA (14–19), HIV-1 reverse transcripts (6) and DNA:RNA R-loops (13). DNA:RNA hybrids are produced during reverse transcription (35), and also DNA repair (36) and transcription (37). We thus hypothesized that as RPRD2 negatively regulates cellular transcription (10), it may also regulate transcription of nascent HIV-1 transcripts, potentially via the same nucleic acid binding mechanism.

This study provides a structural characterization of the RNA-binding properties of RPRD2 by integrating experimental evidence with modern computational prediction methods. Independent analyses using RBDmap, HydRA, RBP-LM and structural interface mapping consistently identified the CID and CREPT domains as the principal regions involved in RNA interactions. Targeted computational mutagenesis demonstrated that substitution of three conserved positively charged residues (Arg90, Arg130 and Lys304) preserved the overall protein structure while weakening interactions with U1 snRNA. Although overall binding to DNA:RNA hybrids was retained, chain-specific analyses revealed reduced interactions with the RNA strand accompanied by enhanced contacts with the DNA strand, suggesting that these residues contribute preferentially to RNA recognition rather than general nucleic acid association. This finding is consistent with previous work that found that RNase treatment increased binding of RPRD2 to immunoprecipitated poly(A) RNA (6). Collectively, these findings provide structural support for previously proposed RNA-binding regions within RPRD2 and offer new mechanistic insights into how the protein may recognize DNA:RNA hybrid substrates during transcriptional regulation and restriction of HIV-1 reverse transcription.

Since RPRD2 is known to bind RNA polymerase II it was perhaps unsurprising that it localized to SC35-enriched nuclear foci, placing RPRD2 at the site of transcription of both host and viral RNA. Due to the long, intrinsically disordered C terminal of RPRD2, it is tempting to question whether in addition to being localized to nuclear speckles, RPRD2 may have a role in formation and/or maintenance of them since interaction of disordered protein domains is essential for the phase-phase separation (57). Nuclear speckles are of particular importance to HIV-1 which hijacks SRRM2 to enlarge the speckles, stabilizing CPSF6 puncta, and promoting efficient viral replication (68). Notably, the nuclear speckle marker SRRM2 has been identified as the true target of the widely used “SC35” antibody (70) also used in this study.

Our findings, in addition to previous work (20,27–29), imply that RPRD2 is an interactor of both the PAF1 and HUSH complexes, both well characterized as regulators of viral and endogenous retroelement transcription. For example, in J-Lat cells, which harbor a GFP reporter driven by a repressed HIV-1 long terminal repeat (LTR) promoter, both HUSH (22) and PAF1 (88) complex depletion resulted in promoter reactivation and GFP expression. Since the PAF1 complex is also a general regulator of host transcription and is localized to the site of transcription (28,89), the interaction between RPRD2 and PAF1 complex members is likely indicative of their roles in regulating cellular transcription. Our results suggest that RPRD2 binds TASOR and the PAF1 complex independently, perhaps suggesting two distinct roles in the vicinity of transcription, or targeting of distinct transcriptional contexts. The apparent co-dependency of RPRD2 and TASOR indicated by the loss of both RPRD2 and TASOR mRNA when either was depleted with siRNA suggests some feedback mechanism between transcription of the two genes. Conversely, we also observed a decrease of TASOR protein when RPRD2 was overexpressed, possibly suggesting that RPRD2 regulates expression of TASOR either directly by affecting TASOR transcription or indirectly via a negative-feedback loop. Regardless of mechanism, since both mRNAs are depleted in both knockdown conditions, it makes interpretation of the phenotype difficult to assign to one protein or the other.

We hypothesized that in addition to inhibiting reverse transcription, RPRD2 may restrict HIV-1 infection by targeting expression of proviral DNA. Given that intron-less DNA is a hallmark of retrovirus and is HUSH-sensitive (26), we predicted enhanced susceptibility of intron-less DNA to RPRD2-mediated transcriptional repression relative to intron-containing DNA. There is still debate as to whether HUSH regulates viral transcription as several studies have shown that the HUSH complex functions to repress transcription of HIV-1 DNA (62–64), but other studies showed that HUSH had no effect on HIV-1 transcription (90–93).

Our initial findings using transfected plasmid-borne LTR-reporter constructs suggested that RPRD2 negatively regulates LTR-driven transcription. The lack of additive effect when both RPRD2 and TASOR were depleted further suggested that RPRD2 may be working via the same mechanism as TASOR, potentially as part of the HUSH complex. In contrast, overexpression of either RPRD2 or TASOR did not significantly alter the percentage of GFP positive cells, consistent with the mechanism of a protein complex, where increased abundance of one subunit does not necessarily increase abundance or activity of the entire complex. Despite these findings, when we expanded our studies to investigate transcription of integrated LTR-driven transcription, we repeatedly observed no impact of RPRD2 depletion. We conclude that RPRD2 does not inhibit transcription of integrated HIV-1 DNA but does restrict transcription of transfected plasmid DNA. It remains possible that RPRD2 affects episomal HIV-1 DNA only, such as 1-LTR and 2-LTR circles, as is seen for other HUSH-related and HUSH-independent cellular factors, such as in the NP220 mediated repression of transcription from murine leukemia virus episomes (90). Given that episomal DNA does not contribute to HIV-1 infection, it is unlikely that this transcriptional inhibition would contribute to the antiviral effect of RPRD2. Further investigation of the mechanisms regulating transcription of unintegrated versus integrated DNA, including the involvement of RPRD2, could potentially be pertinent for understanding the suppression of viral-derived episomal DNA, including in the context of gene therapy.

Lack of effect on endogenous retroelement transcription is surprising given that RPRD2 was identified as a strongly enriched interactor of the HUSH component TASOR, even above the two other recognized members of the HUSH complex MPP8 and Perphilin (20), strongly suggesting it is bound to the complex and involved in its regulatory activity. Based on our findings, it seems likely that either RPRD2 is redundant or dispensable to the HUSH complex activity, or that its interaction is illustrative of the close proximity of RPRD2 and HUSH during transcription. Furthermore, RPRD2 was identified as a regulator of LINE-1 in a genome-wide screen (25). The role of RPRD2 in regulating LINE-1 transcription could be context-dependent; however, we used several cell types, including the K562 line used in the screen. It is relevant that the reporter assay used in that study included a reverse transcription step during retrotransposition, so this could explain why RPRD2 was identified as a hit, in contrast to the results we observed based on transcription alone. RPRD2 has been found by us and others to bind TASOR (20) however neither experiment determined whether RPRD2 was bound to TASOR as part of the HUSH complex. It is possible that RPRD2 binds to TASOR in a HUSH-independent manner. Several distinct functions have been proposed for TASOR outside the canonical HUSH-mediated heterochromatin pathway including as a co-transcriptional RNA quality-control surveillance factor (64).

RPRD2 has been shown to not be an ISG (9), however, RPRD2 mRNA was decreased within 3 hours of IFN type I or 6 hours of type II treatment (85,86), suggesting a possible feedback mechanism. In addition to triggering expression of ISGs, IFN signaling activates the JAK-STAT pathway, leading to expression of repressive transcription factors (94–96). Factors involved in IFN production or signaling are deliberately repressed, once the response is imitated to prevent prolonged or uncontrolled activation (96–99). Since RPRD2 can bind nucleic acids, we hypothesized that RPRD2 could function as an innate immune sensor and that this may underlie its ability to repress virus and retroelements; however, we did not observe any effect on IFN production in RPRD2 knockdown cells in response to a variety of different nucleic acid species. IFN signaling also actively suppresses genes related to cell growth, division, and normal metabolism. Many proteins required for cell cycle progression and division are downregulated, putting the cell into an energy-conserving or growth-inhibited state. Whilst the mechanism is unclear, loss of RPRD2 has been shown to lead to cell cycle arrest (9,10), suggesting it has a role in normal progression, likely through its regulation of cellular transcription.

Finally, we investigated whether RPRD2 has a role in the transcriptional regulation of antiviral genes. IFITM3 is a potent HIV-1 restriction factor typically highly upregulated in response to IFN-α (100,101), and IFITM3 mRNA expression is significantly reduced in response to IFN when cells are depleted of the PAF1 complex (59). Given the interaction between RPRD2 and the PAF1 complex, we therefore chose IFITM3 as a representative ISG to measure in RPRD2 knockdown cells. Although we saw no effect on IFITM3 mRNA expression, this does not rule out the potential role of RPRD2 in regulating transcription of other immune genes.

It is worth noting that RPRD2 depletion was variable across cell types and targeting methods, and smaller decreases in RPRD2 levels may be insufficient to observe a knockdown phenotype. Additionally, the treatment of cells with exogenous DNA, lentiviral constructs, and nucleic acid agonists may influence RPRD2 mRNA expression, which could mitigate the effects of knockdown targeting. Indeed, treatment of THP-1 cells with poly(I:C) led to an increase in RPRD2 protein level (9). However, we saw no change in RPRD2 mRNA expression upon treatment of control cells with 3p-hpRNA.

In conclusion, despite various evidence suggesting that RPRD2 could regulate transcription of HIV-1 provirus or endogenous retroelements, we did not observe this activity. It instead appears that although RPRD2 controls global host transcription, it is not a regulator of LTR-driven transcription. These findings highlight the different mechanisms of regulation of transcription that involve different factors dependent on DNA sequence and context. Given the clear ability of RPRD2 to inhibit HIV infection and LINE-1 retrotransposition, and that it binds nucleic acid species including RNA:DNA hybrids, future endeavors should instead focus on its ability to disrupt reverse transcription as the likely principle mechanism.

---

**Supplemental Figure 1 | Mutation of high likelihood residues in RPRD2 RNA binding domains does not destabilize the protein**

**A)** Structural impact of mutation on RPRD2 protein and nucleic acid-bound complexes. Predicted structural confidence profiles (pLDDT) of wild-type and mutant RPRD2 across the full-length protein, **B)** structured domains, and **C)** CID domain show comparable confidence distributions, indicating preservation of the overall fold. **D)** Interface-level confidence was assessed using mean predicted aligned error (PAE) and **E)** global structural confidence using predicted TM-score (pTM) across DNA/RNA hybrid- and U1 snRNA-bound RPRD2 models. Comparable confidence metrics between wild-type and mutant structures suggest that the mutation does not substantially alter the predicted global architecture or nucleic acid-binding interface.

**Supplemental Figure 2 | RPRD2 does not regulate IFN secretion in response to nucleic acid**

**A)** Effect of RPRD2 on IFN type I production in response to nucleic acid agonists. Inducible shRNA THP-1 cells were subsequently transfected with increasing doses of 3p-hpRNA, **B)** increasing doses of polyI:C, or **C)** standard, low molecular weight and high molecular weight polyI:C. Transfection reagent alone or untreated controls were used in all experiments. After 24 hours, supernatant was collected from the cells and the type I IFN secretion levels quantified using the HEK Blue reporter system (Invivogen).

**Table S1 | List of antibodies, plasmids, primers, siRNAs and shRNAs**

**Table S2 | Residues at predicted intermolecular interfaces between RPRD2 and a DNA:RNA hybrid, or the U1 snRNA stem-loop 4**

## AUTHOR CONTRIBUTIONS

Conceptualization: K.A.J.-J., R.D.S.; Data Curation: K.A.J.-J., R.A., R.D.S.; Formal Analysis: K.A.J.-J., H.X.L., A.T., R.A., K.D., R.M.F., C.H., C.F.; Funding Acquisition: K.A.J.-J., R.D.S.; Investigation: K.A.J.-J., H.X.L., A.T., R.A., K.D., R.M.F., C.H., C.F.; Methodology: K.A.J.-J., R.A., A.C., R.D.S; Project Administration: R.D.S.; Resources: R.D.S., A.C.; Supervision: K.A.J.-J., R.D.S., A.C.; Validation: K.A.J.-J., A.C. & R.D.S; Visualization: K.A.J.-J., R.A., H.X.L., A.T., K.D.; Writing - Original Draft Preparation: K.A.J.-J., R.A., R.D.S.; Writing - Review & Editing: K.A.J.-J., R.D.S.

## ACKNOWLEDGMENTS

All flow cytometry was carried out with support from the Queen’s Medical Research Institute (QMRI) Flow Cytometry and Cell Sorting Facility, University of Edinburgh. This research was supported by a Medical Research Council Transition Fellowship (MR/N013166/1, K.A.J.-J.), NIH/NIAID funding for the HIV Accessory & Regulatory Complexes (HARC) Center (U54-AI170792, K.A.J.-J.), and a Science Olympiad Foundation grant for medical research (K.A.J.-J.). R.A. was supported by an Overseas Scholarship for PhD in Selected Fields Phase-III Batch-3 from the Higher Education Commission of Pakistan. The funders had no role in the study design, data collection and analysis, decision to publish, or preparation of the manuscript.

## MATERIALS AND METHODS

### Protein structure and RNA-binding domain predictions

The full-length human RPRD2 (UniProt accession Q5VT52) consisting of 1,461 amino acids was used as the primary protein sequence. To investigate the contribution of the experimentally predicted RNA-binding region, a second construct comprising residues 1–335 was generated from the full-length sequence, encompassing both the CREPT and CTD-interacting domain (CID). In addition, the experimentally resolved CID structure corresponding to PDB ID: 4FLB (residues 1–161) was downloaded from the Protein Data Bank and analyzed independently. The RNA sequence and structure of U1 snRNA stem-loop structures were obtained from the Protein Data Bank (PDB ID: 7P0V). An experimentally determined DNA:RNA hybrid structure was downloaded from PDB ID: 2DQP, representing a DNA:RNA duplex similar to those recognized during reverse transcription and cellular transcription. Where high-resolution experimental structures were unavailable, three-dimensional structures were predicted using the AlphaFold Server with default parameters. Default settings were retained for all predictions, including the random seed generation and inference parameters. Prediction quality was assessed using the standard AlphaFold confidence metrics, including the predicted Local Distance Difference Test (pLDDT) for residue-level confidence, Predicted Aligned Error (PAE) for relative domain positioning, predicted Template Modelling (pTM) score for overall structural confidence, and interface predicted Template Modelling (ipTM) score for predicted intermolecular interface confidence.

Protein-nucleic acid interaction complexes were predicted using the AlphaFold Server. Each of the three RPRD2 constructs was modelled independently in complex with either U1 snRNA or the DNA:RNA hybrid structure. For every complex prediction, AlphaFold generated five independent structural models, which were subsequently used for comparative analysis. Interface confidence between interacting molecules was evaluated using the reported ipTM scores, while overall model quality was assessed using pTM, pLDDT and PAE values. Mean ipTM values across the five predicted models were calculated and used for comparative analysis between wild-type and mutant complexes. Predicted structures were visualized using UCSF ChimeraX. Intermolecular interface residues were identified using the ChimeraX contacts command. Protein and nucleic acid residues separated by a maximum distance of 4 Å were considered interacting residues. Contact analyses were performed for all predicted complexes, and residue-level interaction data were exported for downstream analysis (**Table S1**).

Residues consistently observed across multiple interfaces were further examined for their physicochemical properties, interaction frequency, and contribution to electrostatic interactions. To investigate the functional importance of the predicted RNA-binding interface, targeted mutations were introduced into all three RPRD2 constructs at residues Arg90, Arg130, and Lys304. Each residue was individually substituted with glutamic acid (R90E, R130E and K304E) to reverse the local electrostatic charge while maintaining minimal changes to residue size. Three-dimensional structures were predicted for all mutant proteins using the AlphaFold Server under the same prediction settings used for the wild-type proteins. Predicted pLDDT confidence scores were extracted and compared with their corresponding wild-type proteins to determine whether the introduced mutations altered the overall structural integrity of the proteins. Mutation sites were mapped onto residue confidence plots to visualize their structural context. All mutant proteins were subsequently modelled in complex with both U1 snRNA and the DNA:RNA hybrid using the same AlphaFold workflow applied to the wild-type proteins. For each complex, mean ipTM scores from the five predicted models were calculated to assess changes in interface confidence following mutation. PAE values were additionally examined to evaluate the confidence of the predicted intermolecular interfaces. Comparisons between wild-type and mutant complexes were used to determine whether disruption of candidate interface residues altered predicted nucleic acid binding. Predicted wild-type and mutant complexes were further analyzed using the PDBePISA server to characterize intermolecular interfaces and estimate interaction stability. For each complex, PDBePISA was used to calculate interface area, buried and solvent-accessible surface areas, hydrogen bonds, salt bridges, and solvation free-energy gain upon interface formation (ΔiG). These parameters were compared between wild-type and mutant complexes to determine whether mutations altered interface stability or reduced favorable intermolecular interactions. All downstream analyses, statistical summaries and figure generation were performed using Python. Comparative visualizations of AlphaFold confidence metrics, interface properties and PDBePISA-derived parameters were generated to facilitate comparison between protein constructs, nucleic acid ligands and mutant variants.

### Cell lines and cell culture

Human embryonic kidney (HEK) 293T cells (ATCC, CRL-3216) and HeLa cells were generous gifts from Jurgen Haas and were maintained in Dulbecco’s modified Eagle’s medium (DMEM) (Sigma-Aldrich, #D6429). Human THP-1 cells (a generous gift from Sam Wilson) and Jurkat cells (a generous gift from Jurgen Haas) were maintained in Roswell Park Memorial Institute (RPMI) 1640 medium (Gibco, #11875093). J-Lat cell lines (clones A1 and H2) (71) were obtained from the National Institute for Biological Standards and Control (NIBSC) and maintained in RPMI 1640 medium. K562 cells were kindly donated by Dr Conor Poland (Macomics) and maintained in RPMI 1640 medium. All media was supplemented with 10% heat-inactivated fetal bovine serum (HI-FBS) (Gibco, #10082147) and 1% penicillin-streptomycin (50 mg/ml; Corning, #30-001-CI). Cells were cultured at 37°C in a humidified incubator with 5% CO_2_. For J-Lat reactivation experiments, cells were treated with 10 ng/ml TNFα (Sigma-Aldrich, # H8916-10UG) for 24 h to activate transcription from the HIV-1 LTR through the NF-κB pathway (102). Cells were regularly tested for mycoplasma using LookOut Mycoplasma PCR Detection Kit (Sigma Aldrich, #MP0035-1KT).

J-Lat clone A1 and H2 cells, THP-1 cells, and HeLa cells stably expressing inducible short hairpin RNA (shRNA) cassettes against RPRD2 (shRPRD2) or non-targeting control shRNA (shScr) were created as previously described (8,9). The isopropyl-β-D-1-thiogalactopyranoside (IPTG)-inducible vector pLKO-IPTG-3xLacO (Sigma) was used to express shRNAs targeted against RPRD2 (Mission TRCN0000141116; Sigma), or a nontarget (scrambled) control shRNA. Viral particles for cell line transductions were prepared by transfecting HEK293T cells with a mix of 500ng packaging plasmid pCMVΔR8.91, 450ng vesicular stomatitis virus (VSV) G protein plasmid pMD2.G (Addgene, #12259), and 750ng shRPRD2 or shScr pLKO-IPTG-3xLacO plasmid with 5 µL transfection reagent FuGENE HD (Promega). After 48 h, the pseudotype medium was collected, filtered through a 0.45 μm pore-size filter and stored at -80°C as single-use aliquots. J-Lat cells, THP-1 cells and HeLa cells were transduced with viral particles in the presence of 8 g/ml Polybrene, by centrifugation for 90 min at 1500 rpm, incubation overnight at 37°C, followed by replacement of medium. After 48 h, resistant cells were selected using 3 ug/ml puromycin (Gibco, #A1113803) and cultured until all mock-treated negative control cells had died. Surviving cells were maintained in medium supplemented with 0.5 ug/ml puromycin. Culturing cells in the presence of 1 mM IPTG for 72 h induced expression of shRNAs.

### Transfection of expression plasmids and siRNAs

Plasmids expressing RPRD2 with either a GFP or myc tag at the N-terminus (6) were generous gifts from Aine McKnight. Plasmid expressing catalytic mutant RNase H with a GFP tag at the N-terminus was a generous gift from Lee Zou. Plasmid expressing TASOR with a myc tag at the N-terminus was a generous gift from Roy Matkovic. Empty vector plasmids pcDNA3.1(+) eGFP (#129020) and pcDNA3.1 myc (#176045) were obtained from Addgene. The HIV-1 LTR-Tat-IRES-GFP construct (71), the HIV-1 LTR-driven intron-containing and intron-less GFP reporter constructs (pHRSIN-as(PSFFV-GFP*-(-1)-ATGless-ORF2-SV40pA)-pSV40-Blast and pHRSIN-as(PSFFV-GFP*-(-1)-ATGless-ORF2-(5’HBB IVS2)-SV40pA)-pSV40-Blast) (26), and the L1-derived intron-containing and intron-less iRFP piggyBac reporter constructs (26) were all generous gifts from Paul Lehner. Predesigned siRNA pools targeting RPRD2 and TASOR were purchased as TriFECTa kits containing 3 target-specific Dicer-Substrate siRNAs (Integrated DNA Technologies (IDT)). Specific siRNAs targeting RPRD2 (CACGTAAGCCCTCAGATGATA), TASOR (SASI_Hs02_00325516) or non-targeting control were obtained from Sigma Aldrich. The shRNA plasmid used for transient knockdown of RPRD2 was pSUPER.retro.puro-RPRD2, created using the pSUPER RNAi system (OligoEngine) by adding the RPRD2 specific insert sequence: 5′ AGATCCCCCC**CACGTAAGCCCTCAGATGA**<u>TTCAAGAGA</u>TCATCTGAGGGCTTACGTGTTTTTAAGCT T 3’, where the target sequence is in bold and the hairpin loop is underlines, into the pSUPER.retro.puro base vector.

Plasmids and siRNAs were transfected into HEK293T cells using Lipofectamine 2000 reagent (Invitrogen) or polyethylenimine (PEI) (Polysciences, 23966) at a 1:3 ratio of DNA or RNA to transfection reagent, according to manufacturer’s protocol. Medium was replaced 6 h after transfection to reduce cytotoxicity. For co-transfection, 125 ng pLTR-Tat-IRES-GFP and 10 pmol (20 nM) siRNA were mixed with 0.5 µL lipofectamine 2000 in 25 µL Opti-MEM reduced serum medium (Gibco, #31985062) per well of a 24-well plate. For double knockdown (DKD) of both RPRD2 and TASOR 10 pmol of each siRNA was transfected. Inducible-shRNA cells were transfected after 24 h of IPTG treatment. After 48 h cells were subject to either RNA isolation for qRT-PCR or flow cytometry analysis of GFP reporter expression.

### Immunoprecipitation

Cells were washed with ice-cold phosphate-buffered saline (PBS), then lysed in ice-cold immunoprecipitation (IP) buffer [20 mM Tris-HCl pH 7.5, 150 mM NaCl, 1mM EDTA, 1% Nonidet P40 Substitute, and 0.2% sodium deoxycholate] freshly supplemented with PhosphoSTOP phosphatase inhibitor tablets (Roche) and COMplete Protease Inhibitor tablets (Roche) for 20 min on ice. Chromatin was sheared by sonification (30 sec x 10 cycles, amplitude 5) then samples were centrifuged at 13,300 rpm for 30 min, and supernatant transferred to a fresh tube. Protein concentration was quantified and normalized with the bicinchoninic acid assay (BCA) (Pierce BCA Protein Assay Kit, Thermo Scientific) according to manufacturer’s protocol.

Lysates were precleared with Dynabeads Protein-G (Novex, Life Technologies) for 1 h, rotating at 4°C. Anti-DNA:RNA hybrid (S9.6) antibody (56), or antibodies recognizing RPRD2 (Abcam, ab83823), PAF1 (Abcam, ab70638) or TASOR (Abcam, ab322400) were conjugated to magnetic beads (Sigma-Aldrich, #M8823) and washed 3 times with IP buffer before incubation with the precleared lysate overnight rotating at 4°C. Beads were washed 5 times, each for 5 min with IP buffer and then bound protein was eluted by boiling in SDS sample buffer supplemented with reducing agent for 5 min. All antibodies used can be found in **Table S1**. For DNA:RNA hybrid immunoprecipitation, cells were transfected with RPRD2-GFP or mutant RNase H-GFP plasmid 48 h prior to harvest, as described above, except cells were crosslinked with UV and lysed with Chromatin Immunoprecipitation Sequencing chromatin immunoprecipitation (ChIP) buffer (SimpleChIP Sonication Cell and Nuclear Lysis Buffer, Cell Signaling, #81804).

### Protein isolation and immunoblotting

At 48 h post-transfection, cells were washed with phosphate-buffered saline (PBS), then lysed in radioimmunoprecipitation assay (RIPA) buffer [20 mM Tris-HCl, 150 mM NaCl, 1 % Triton, 1% sodium dodecyl sulphate]. RIPA buffer was freshly supplemented with phosphatase inhibitor tablets (Roche), protease inhibitor tablets (Roche), 3 mM Benzonase (Merck,) and 5 mM MgCl. Lysates were centrifuged at 17,000 × g for 20 min, supernatant moved to a fresh tube and stored at –20°C. Protein concentration were determined with Pierce™ BCA Protein Assay Kit (ThermoFisher, #23227). Lysates were boiled with NuPAGE™ LDS (lithium dodecyl sulfate) Sample Buffer (4X) (ThermoFisher, # NP0007) and with 50 mM dithiothreitol (DTT) for 5 min at 95°C, then stored at –20°C.

Samples were thawed and for each sample 30-50 ug of protein was loaded onto NuPAGE 4-12% Bis-Tris, 1.0 mm precast SDS-PAGE protein gels (ThermoFisher) in addition to 5 μL of PageRuler Plus prestained protein ladder (Thermo Scientific, #26619). Proteins were resolved by SDS-PAGE (NuPAGE Protein gels, Invitrogen) at 130 V for 90 min in MOPS running buffer (ThermoFisher). Proteins were transferred to 0.45 μm pore-size PVDF membrane in MOPS transfer buffer (ThermoFisher) by methanol-based electrotransfer using an XCell II blot module (Invitrogen) at 90 V for 2 h. Membranes were stained with Ponceau S and then blocked in 5% (w/v) milk powder in tris-buffered saline with 0.1% Tween 20 (TBS-T) for 30 min at room temperature. Membranes were cut accordingly and incubated with rabbit anti-DNA:RNA hybrid (S9.6) (1:1000) (Sigma-Aldrich, #MABE1095), rabbit anti-RPRD2 (1:1000) (Abcam, ab83823), rabbit anti-TASOR (1:1000) (Abcam, ab322400), or rabbit anti-GAPDH (1:5000) (Abcam, # ab181602) primary antibody diluted in 5% (w/v) bovine serum albumin (BSA) in TBS-T at 4°C overnight. After washing, membranes were incubated with goat anti-rabbit immunoglobulin (IgG) horseradish peroxidase (HRP)-conjugated secondary antibody (1:10,000) (Abcam, ab97051) for 1 h at room temperature. Signal was detected using SuperSignal West Pico developer (ThermoFisher) for 5 min and imaged on Hyblot ES Autoradiography Film (Denville Scientific). Blots were incubated in 1X ReBlot Plus Antibody Stripping Solution (Millipore) before reprobing.

### Confocal immunofluorescence microscopy

HEK293T cells were grown on coverslips and transfected with RPRD2-GFP plasmid as described above. After 48 hr, cells were fixed with 4% paraformaldehyde at room temperature for 10 min. Coverslips were then washed 3 times with PBS, stained with 4,6-diaminidino-2phenylidole (DAPI) at 50ng/ml, mounted in Vectashield (Vector), sealed with nail varnish, and cured in the dark overnight. For nuclear speckle imaging, after fixation cells were permeabilized with the Cytofix/Cytoperm kit (BD Biosciences) / 0.5% Triton X-100 at room temperature for 10 min. Coverslips were then incubated for 1 h with block buffer [1% BSA, 0.01% Triton X-100 in PBS] at room temperature, followed by incubation with rabbit anti-SC35 primary antibody (diluted 1:500 in block buffer) in a humidified chamber overnight at 4°C. Coverslips were washed 3 times with wash buffer (0.01% Triton X-100 in PBS), and incubated with Alexa Fluor 594 secondary antibody (Molecular Probes, diluted 1:1000 in block buffer) in a dark, humidified chamber for 1 h at room temperature. Coverslips were then washed 3 times with wash buffer, followed by 3 PBS washes, DAPI staining and mounting as described above. Data were acquired using a Leica TCS SP5 confocal microscope at 20x magnification and NIS Elements AR software (Nikon Instruments Europe, Netherlands). Z-stacks of images were acquired with a 0.2 µm step, scan size 1024x1024, 1.2x zoom and 2x frame averaging. Image analysis was carried out using the FIJI/ImageJ software.

### RNA extraction and quantitative reverse transcription polymerase chain reaction (qRT-PCR)

Total RNA was extracted from cells using ReliaPrep RNA Miniprep Systems (Promega) according to the manufacturer’s instructions. Reverse transcription quantitative polymerase chain reaction (qRT-PCR) was performed using Brilliant II SYBR Master Mix one-step RT–PCR kit (Agilent) following the manufacturer’s instructions and run on a LightCycler 96 Instrument real-time PCR system (Roche) following this program: reverse transcription at 50°C for 30 min, 95°C for 2 min, then 40 cycles of 95°C for 30 sec, 55°C for 20 sec, 70°C for 20 sec followed by the plate read step. Each sample was performed in technical triplicate. Primers used were: RPRD2 (Forward – GAATCATTTGCTGATGTGCTTCC, Reverse – CTCTCAATGCCACAATCATGTCT), TASOR (Forward – TGAAGACATTGCAGGTTTCATTC, Reverse – CATCCAGGCTATCAACACCAG), LINE-1 5′UTR (#1) (Forward – GAATGATTTTGACGAGCTGAGAGAA, Reverse – GTCCTCCCGTAGCTCAGAGTAATT), LINE-1 5′UTR (#2) (Forward – ACAGCTTTGAAGAGAGCAGTGGTT, Reverse – AGTCTGCCCGTTCTCAGATCT), HERV-K gag (Forward – AGCAGGTCAGGTGCCTGTAACATT, Reverse – TGGTGCCGTAGGATTAAGTCTCCT) and β-actin (Forward – CGCGAGAGAAGATGACCCAGATC, Reverse – GCCAGAGGCGTACAGGGATA). LightCycler 96 software v1.1 was used for analysis. Gene expression data was analyzed by the delta delta Ct method. Expression level of RPRD2 and TASOR were normalized to expression level of β-actin housekeeping control. Data are presented as fold change compared to negative control. Unpaired two-tailed t-tests were used for statistical analysis.

### Flow cytometry and data analysis

Flow cytometry analysis was performed on an Attune NxT Flow Cytometer (Thermo Fisher Scientific), recording all events in a 50 μL sample volume after one 150 μL mixing cycle. Data were exported as FCS3.0 files using Attune NxT Software v3.2.0 and analyzed with a consistent template on FlowJo™ Software (BD Biosciences) (**Figure S1**). Cells were gated for lymphocytes by light scatter followed by doublet discrimination in both side and forward scatter. Cells with equal fluorescence in the BL-1 (GFP) channel and the VL-1 channel were identified as autofluorescent and excluded from the analysis. A consistent gate was then used to quantify the fraction of remaining cells that expressed GFP. All flow cytometry was carried out with support from the Queen’s Medical Research Institute (QMRI) Flow Cytometry and Cell Sorting Facility, University of Edinburgh.

### Lentiviral based pseudotype production and transduction

HIV-1-derived LTR-driven intron-less and intron-containing GFP reporter plasmids were kind gifts from Paul Lehner (26). Reporters were delivered to cells by lentiviral transduction. HEK293T cells were co-transfected with packaging plasmid pCMVΔR8.91, vesicular stomatitis virus (VSV) G protein plasmid pMD2.G (Addgene, #12259) and the relevant reporter plasmid using polyethylenimine (PEI) (Polysciences, 23966). Supernatant was harvested at 48 h post-transfection, passed through a 0.45 μm filter, and stored at -80°C. Target cells were seeded at 6x10^4^ cells/well in 48-well plates. Reduced serum medium, Opti-MEM (ThermoFisher), was used to dilute viral supernatant and increase the efficiency of transduction. Transduction was performed via spinoculation at 1800 rpm for 1 h with a total volume of 100 µL viral dilution, followed by incubation for 3 h prior to adding maintenance medium. After 48 h incubation, transduced cells were selected with Blasticidin (5μg/ml).

### PiggyBac transposon integration and expression

L1-derived piggyBac intron-less and intron-containing iRFP reporter plasmids and the piggyBac transposase-expression plasmid were kind gifts from Paul Lehner (26). HeLa cells were cultured in DMEM supplemented with Tet-free HI-FBS (Qualified One Shot, ThermoFisher). Co-transfection of each iRFP reporter plasmid and piggyBac transposase plasmid at ratio 5:1 was performed using FuGENE (Promega), according to manufacturer’s instructions. After 48 h, transfected cells were selected with Blasticidin (5μg/ml) for at least 7 days. Expression of the piggyBac intron-less or intron-containing iRFP reporters was induced by doxycycline (1 μg/ml) treatment, and 24 h later expression was measured by flow cytometry.

### CRISPR-Cas9 genome editing

Two crRNA sequences targeting exon 1 of RPRD2; crRNA #1 (CCAGTCGGTAACCAACACCA), crRNA #2 (CCATGGTGTTGGTTACCGAC), and a crRNA sequence targeting IFITM1 (UGAUCACGGUGGA) were ordered as part of the Alt-R™ CRISPR-Cas9 System (Integrated DNA Technologies). Ribonucleoprotein (RNP) complexes were prepared by mixing 1 μL of both crRNA and tracrRNA were combined in a 1:1 M ratio (1 μM) in 98 μL of Alt-R™ Suspension Buffer, heated at 95°C for 5 min and cooled to room temperature to form crRNA:tracrRNA gRNA. 38.75 μL of the gRNA suspension (1 μM) was combined with an equal volume of Cas9 suspended in Opti-MEM Reduced Serum Medium (1 μM) (ThermoFisher). 35 μL Opti-MEM was added to the gRNA-Cas9 suspension alongside 12.5 μL of CRISPRMAX™ Cas9 Plus Transfection Reagent (ThermoFisher) and incubated at room temperature for 5 min. 7.5 μL of CRISPRMAX™ Reagent (ThermoFisher) was mixed with 117.5 μL Opti-MEM to form a separate suspension. Both suspensions were combined and incubated for 15 min at room temperature. K562 cells were diluted to 200,000 cells/mL in antibiotic-free RPMI, and 2.25 mL was added to each well of a 6-well plate. 250 μL of the RNP complex suspension was added to each well and incubated for 48 h at 37°C prior to protein isolation for western blot analysis and RNA isolation for qRT-PCR analysis.

### Data from publicly available databases

Constitutive expression of RPRD2 mRNA and protein expression was accessed from The Human Protein Atlas database of tissue expression (81,82). Immune cell expression was accessed from the Monaco dataset (83). To investigate RPRD2 change in expression in response to cytokines we accessed the Immune Dictionary (84). RPRD2 expression changes in response to IFN was accessed from the Interferome: the IFN database (85) and the Orthologous Clusters of Interferon-Stimulated Genes database (86). The STRING database (https://string-db.org/) (103,104) was employed to retrieve experimentally confirmed and predicted protein-protein interaction and generate a visual protein interaction network.

### Interferon (IFN) Secretion Assay

The HEK Blue reporter system (InvivoGen) was used to assay for type I interferon (IFN) production. Activation of IFN signaling leads to expression of a secreted embryonic alkaline phosphatase (SEAP) reporter gene, which can be detected and its absorbance measured via spectrophotometry. Inducible shRNA THP-1 cells were created as described above and treated with IPTG to induce knockdown. 24 h later, cells were treated with polyI:C (0.1, 1, 10 or 25 μg/mL, InvivoGen), 3p-hpRNA (64, 32 or 16 ng/mL, InvivoGen), polydA:dT (500 ng/mL, InvivoGen), or polydG:dC (500 ng/mL, InvivoGen), 25 μg/mL HMW poly(I:C) (Invivogen), or 25 μg/mL LMW poly(I:C) (InvivoGen) using Lipofectamine 2000 (Invitrogen) as per manufacturer’s protocol, mock transfected or untreated. After 24 h, culture media from the THP-1 cells was added to the HEK Blue cells to activate the reporter gene. In addition, an IFN-β standard curve was prepared at the following concentrations (in ng/mL): 35, 25, 17.5, 12.5, 10, 8, 6, 4, 2, 0.5, 0.25, diluted from a stock solution of 100ng/mL IFN-β in PBS, and also added to HEK Blue cells. After 24 h cell culture media was mixed with QUANTI-Blue solution (Invivogen), absorbance quantified using a CLARIOstar microplate reader (BMG Labtech), and sample values compared to the standard curve to quantify IFN secretion.

### Quantification and statistical analysis

Data were analyzed using GraphPad Prism 10 (GraphPad Software, La Jolla, CA). Error bars represent mean ± standard deviation unless otherwise stated. For all figures: * = p<0.05, ** = p<0.01, *** = p<0.001, **** = p<0.0001. Every experiment included at least three technical replicates. Statistical significance in all experiments was calculated by analysis of variance (ANOVA).

### Data and code availability

This paper does not report novel datasets or original code. Subfigures and plots were generated using GraphPad Prism 10 (GraphPad Software, La Jolla, CA). All figures were assembled in Adobe Illustrator (Adobe, Inc.).

