## Supplementary figures and images for "The HIV-1 restriction factor RPRD2 does not inhibit transcription of HIV-1 or endogenous retroelements"

### Supplemental Figures

#### Supplemental Figure 1

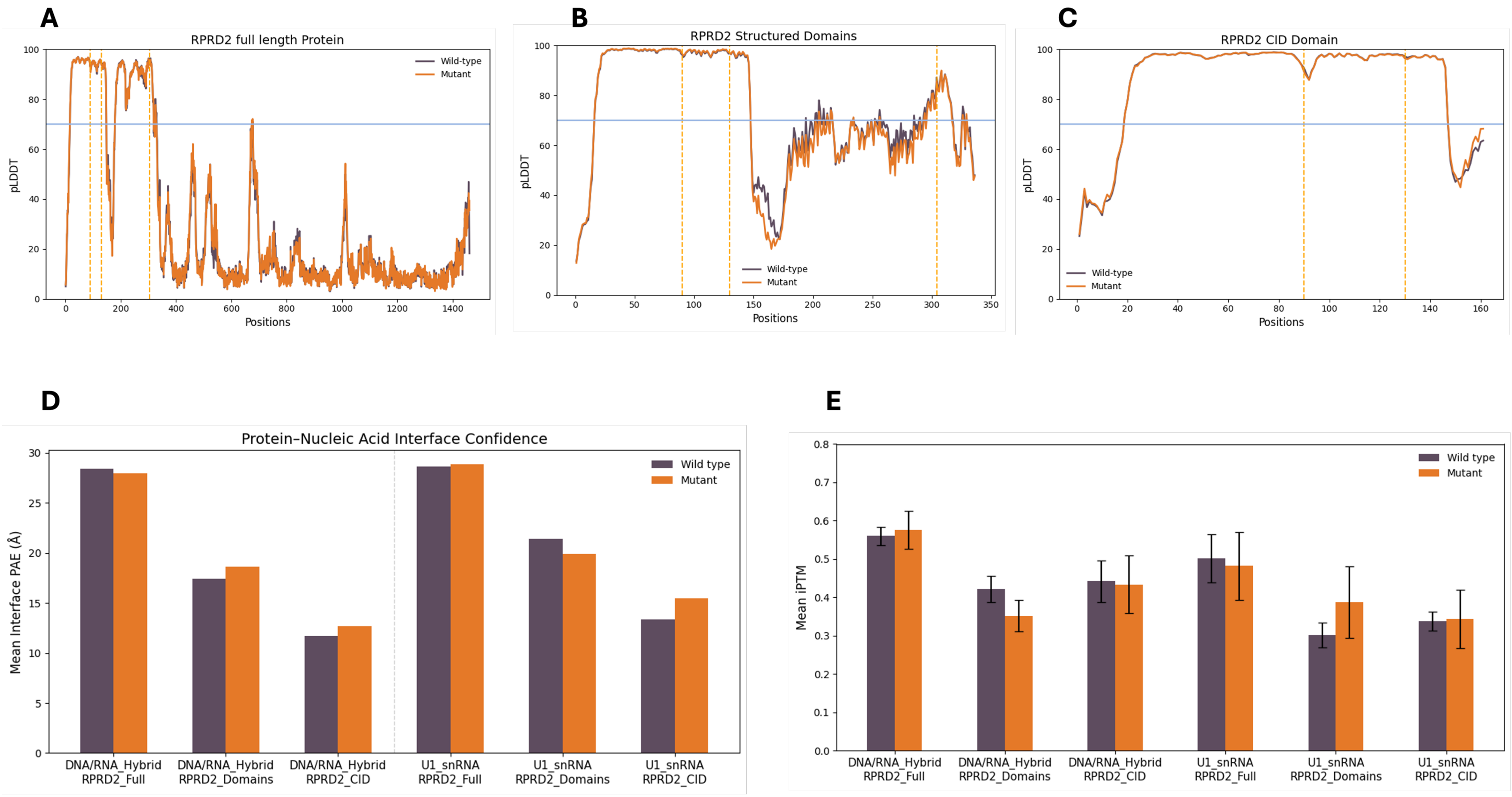

#### Supplemental Figure 2

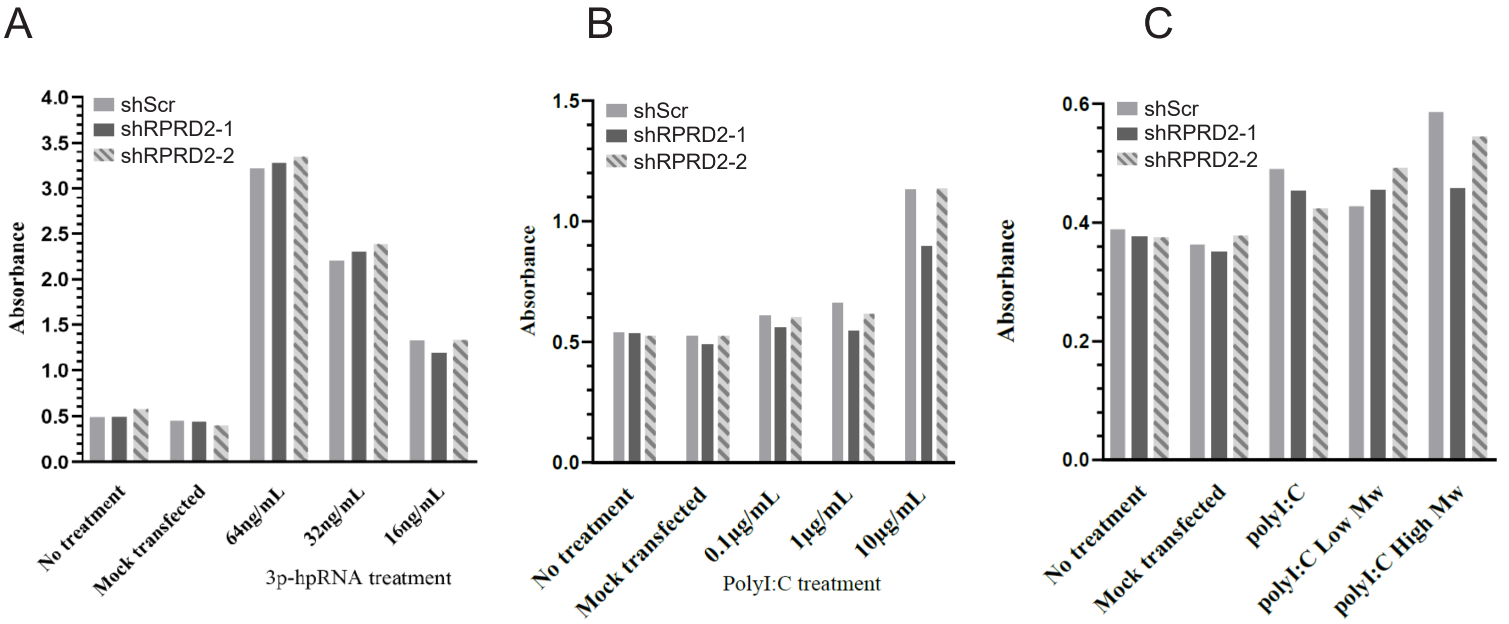
